# Quantifying and contextualizing the impact of bioRxiv preprints through automated social media audience segmentation

**DOI:** 10.1101/2020.03.06.981589

**Authors:** Jedidiah Carlson, Kelley Harris

**Affiliations:** Department of Genome Sciences, University of Washington, Seattle, WA; Computational Biology Division, Fred Hutchinson Cancer Research Center, Seattle, WA

## Abstract

Engagement with scientific manuscripts is frequently facilitated by Twitter and other social media platforms. As such, the demographics of a paper’s social media audience provide a wealth of information about how scholarly research is transmitted, consumed, and interpreted by online communities. By paying attention to public perceptions of their publications, scientists can learn whether their research is stimulating positive scholarly and public thought. They can also become aware of potentially negative patterns of interest from groups that misinterpret their work in harmful ways, either willfully or unintentionally, and devise strategies for altering their messaging to mitigate these impacts. In this study, we collected 331,696 Twitter posts referencing 1,800 highly tweeted bioRxiv preprints and leveraged topic modeling to infer the characteristics of various communities engaging with each preprint on Twitter. We agnostically learned the characteristics of these audience sectors from keywords each user’s followers provide in their Twitter biographies. We estimate that 96% of the preprints analyzed are dominated by academic audiences on Twitter, suggesting that social media attention does not always correspond to greater public exposure. We further demonstrate how our audience segmentation method can quantify the level of interest from non-specialist audience sectors such as mental health advocates, dog lovers, video game developers, vegans, bitcoin investors, conspiracy theorists, journalists, religious groups, and political constituencies. Surprisingly, we also found that 10% of the highly tweeted preprints analyzed have sizable (>5%) audience sectors that are associated with right-wing white nationalist communities. Although none of these preprints intentionally espouse any right-wing extremist messages, cases exist where extremist appropriation comprises more than 50% of the tweets referencing a given preprint. These results present unique opportunities for improving and contextualizing research evaluation as well as shedding light on the unavoidable challenges of scientific discourse afforded by social media.

## Introduction

The primary barometer of success in academic research is the extent to which the research (or individual researcher) has contributed to new discoveries and advancement of knowledge. Quantifying, predicting, and evaluating these achievements plays a pivotal role in the academic research ecosystem, and the metrics used to assess the impact of scientific publications can have a profound influence on researchers’ career trajectories. Measuring the scientific impact of a publication typically involves metrics based on the number of citations received in the literature. Examples of such metrics include the h-index, which is the maximum number *h* for which an author has published *h* or more papers with at least *h* citations; and the Journal Impact Factor, which is the average annual number of citations received by articles published in a given journal [1]. These citation-based metrics (hereafter referred to as “traditional scientometrics”) are not without their flaws, as they can be gamed and exploited [2] and are susceptible to systematic biases [3]. Nevertheless, traditional scientometrics provide a standardized and easy-to-interpret system of evaluating the scientific impact made by an individual article or researcher, or even entire institutions or academic journals.

Beyond the intrinsic goal of knowledge production, publicly-funded research is typically expected to produce some form of societal impact in the form of social, cultural, environmental, or economic returns for the taxpayer [4], and researchers are regularly expected to justify how their line of study will generate such impacts [5]. For example, the importance of societal impact is codified in National Science Foundation grant application guidelines, where funding applicants must dedicate a substantial section of their proposals to describing the “potential [for their research] to benefit society and contribute to the achievement of specific, desired societal outcomes” [6]. This distinction between scientific and societal impacts is relatively new: as noted by Bornmann [4], for most of the 20th century, policymakers believed that any investment in scientific research naturally led to positive societal impact, so the only relevant impact to be evaluated was the degree to which a particular research output or individual researcher advanced high-level academic knowledge.

However, a common point of concern is that existing paradigms for evaluating scientific impact are often inappropriate for assessing the societal implications of scientific research [7]. One reason for this discordance is simply the slow pace at which literature citations accumulate: it may take years or even decades for an impactful article to accrue its due attention in the scientific literature, which could lead to the short-term conclusion that the research was neither scientifically nor societally impactful. Moreover, literature citations only directly measure the attention received from other researchers and tell us very little about how a given article of research has penetrated the public mindset. An important goal in research evaluation, therefore, is to identify data sources that qualitatively and quantitatively trace engagement with research from non-specialist audiences.

### Altmetrics: a new way to measure societal impact?

In the last decade, scientists have flocked to Twitter and other online social media platforms to share their research, connect with colleagues, and engage with the public [8]. The enthusiastic adoption of social media by some researchers has dramatically altered how scientific publications are diffused throughout the scientific community and bridged into the public forum, leading some in the research community to consider social media as a vital tool for public engagement and scientific communication [9]. Unlike traditional metrics, data gleaned from social media (and metrics thereof, commonly known as “altmetrics”) can provide a nearly instantaneous readout of a paper’s exposure across broad cross-sections of experts and lay audiences alike. Though some have speculated that “social media buzz” about research has the potential to be highly superficial and not necessarily indicative of lasting societal or scientific impact (or even if the paper is downloaded and read) [10–12], other evidence suggests that a flurry of interest in new research on social media might actually provide an immediate indicator of its potential downstream scientific and societal impacts, including: 1) predicting the number of times one’s research is cited in the scientific literature [13]; 2) facilitating engagement with news media and public audiences [14,15]; 3) fostering networking, collaboration, and career development opportunities [9] and 4) predicting the likelihood of receiving grant funding [16].

In 2010, the “Altmetrics Manifesto” outlined a vision for how social media data and other altmetrics could expand our ability to systematically measure and assess the societal impact of research [17]. In the following years, a cottage industry of organizations, such as Altmetrics, ImpactStory, and Elsevier’s Plum Analytics, have brought the concept of altmetrics to the mainstream. For millions of journal articles, these companies now index every instance in which an article is referenced on social media sites like Twitter, Facebook and Reddit, along with mentions in news articles, Wikipedia pages, and policy documents. These data are typically aggregated into a single article-level score, largely weighted by how frequently the article is referenced on social media. Preprints that garner high altmetric scores are celebrated by some authors, climb to the top of article recommendation engines, and provide prospective readers with a glimpse of which new research appears to be generating attention within days or even hours of its initial public release.

Despite the general enthusiasm about altmetrics, metaresearch experts have stressed that singular quantitative indicators like the Altmetric Attention Score fail to provide the appropriate context necessary for evaluating societal impact [7,18]. Altmetric readily acknowledges that their Attention Score alone is insufficient for evaluating a paper’s scope of impact: In Altmetric’s introduction to the list of papers that received the top 100 Attention Scores of 2019, they caution that “the only theme that many of these papers have in common is their ability to start conversations…the ranking has no bearing on the quality or impact of the research itself.” [19]. Identifying these “conversation starter” papers is certainly a valuable indicator of research that might lead to tangible impacts on the broader public, but if altmetrics are to be harnessed to meaningfully measure societal impact, we must move beyond the simple question of *how much* attention an article receives on social media, and prioritize questions of context: *who* is engaging with the article, *what* they are saying about it, and *when* and *where* they are doing so [7,20].

### Contextualizing altmetrics

Among the major altmetrics data aggregators (Altmetric, Plum Analytics, Impact Story), only Altmetric attempts to provide cursory classifications of Twitter users who have engaged with an article. Altmetric uses a proprietary algorithm that evaluates “keywords in profile descriptions, the types of journals that users link to, and follower lists” from each user to segment the overall audience into four basic categories: three describe different groups of academic/scientific stakeholders (“scientists,” “practitioners,” “science communicators”), and a fourth, “members of the public,” encompasses all others who do not fit into one of the other categories [21]. In a blog post written for Altmetric [22], Haustein, Toupin, and Alperin explain that classifying users even into these coarsely-defined groups is a challenging task due to the inherent sparsity and noisiness of direct self-descriptions provided by Twitter users in their biographies, which are limited to just 160 characters. Moreover, Altmetric’s audience classifications do not appear to factor into how Altmetric Attention Scores are quantified, nor are these data directly visible on publishers’ websites alongside the Altmetric Attention Scores for each article. Despite Altmetric’s repeated emphasis that demographic context is important, these details appear to be downplayed in favor of a singular quantitative score.

Fortunately, we are not restricted to learning about audience characteristics from individual Twitter biographies, which are plagued by sparsity and noise. A cornerstone of sociological research is the principle of network homophily, which states that individuals tend to connect and engage within a social network of others who share similar characteristics [23]. Online social media is no exception: patterns of network homophily have been demonstrated in numerous studies of Twitter users [24–28], and there is evidence to suggest that the homophily of an individual’s online connections generally mirrors their face-to-face networks [29]. Recent studies have applied this principle to the study of altmetrics and demonstrated that deeper contextualization of a publication’s audience on social media can be achieved by examining various aspects of how individual users are networked with others on Twitter [7,30]. More specifically, network homophily enables the identification of various characteristics of an individual Twitter user based on the self-descriptions of the accounts connected with that individual on Twitter [25,31].

In this study, we present a framework for classifying members of a paper’s audience on Twitter into granular, informative categories inferred through probabilistic topic modeling of metadata collected from each user’s network of followers. Probabilistic topic modeling is a powerful and flexible method for revealing the granular composition of all sorts of complex datasets—not only the present application of text mining [36], but also inference of population ancestry using genetic markers [32], computer vision [33], and many other areas of study. With this approach, we analyze the Twitter audiences for 1,800 highly tweeted preprints (encompassing over 330,000 tweets in total) across a variety of topics in biology. We show that each article’s audience on social media is characterized by a unique composition of online communities, both within academia and among diverse lay audiences. The audience classifications inferred with our topic modeling framework thus provide valuable context for interpreting altmetrics and providing initial traces of potential societal impacts.

We highlight three ways in which these inferred audience demographics can serve to enhance interpretation of altmetrics: 1) more accurate quantification and classification of the various academic audience sectors that express interest in a particular research output, 2) detailed characterization of lay audiences that inform potential societal impacts, and 3) exploring how politically-engaged audiences are interacting with research. Because detailed context is such an important aspect of interpreting altmetrics, we have compiled an online portal showcasing summary data across all preprints analyzed along with interactive visualizations detailing the various dimensions of audience characteristics for individual preprints, available at http://carjed.github.io/audiences.

Our analyses also reveal a subset of preprints that attract a great deal of attention from audience sectors associated with far-right ideologies, including white nationalism. These communities appear to be especially active in their engagement with preprints concerning the genetic architecture of behavioral traits, human population genetics and ancient DNA research, and the neurological and physiological variation across sexes and genders, among a plethora of other topics. In some cases, these politically-motivated audience sectors comprise over half of the total audience of users referencing an article, providing concrete evidence to support concerns about racist and sexist misappropriation of research that have been expressed by academic organizations [34], news media [35], and scientists themselves [36]. We discuss how stakeholders in the scientific community can use these audience demographics to understand the social and political implications of their work and assert their expertise to guide public science literacy in the era of social media.

## Results

### Data collection

We used Rxivist, a service that indexes article-level metrics for preprints posted to the bioRxiv preprint repository [37], to identify 1,800 of the most highly tweeted bioRxiv preprints posted between November, 2013 and February, 2020, spanning the 27 categories under which bioRxiv preprints are organized. The number of tweets we analyzed per preprint ranged from a minimum of 50 to a maximum of 3,794 tweets. We focused on bioRxiv preprints rather than peer-reviewed journals for the following reasons: 1) bioRxiv covers a broad cross-section of topics in the biological sciences, 2) metadata for bioRxiv preprints is readily retrievable through the open-source Rxivist platform, 3) bioRxiv preprints are fully open-access and freely available to read by anyone, and 4) unlike many publishers of peer-reviewed research, bioRxiv does not highlight or promote certain preprints over others, nor are preprints typically promoted to the news media via press releases, creating a more even playing field for preprints to organically garner attention on social media.

### Topic modeling for audience sector classification

Given a preprint and the list of Twitter users that tweeted or retweeted a link to it, we are primarily interested in classifying each of these users into a particular “audience sector,” defined here as a subset of users whose followers share similar self-identified characteristics (e.g., occupation, personal interests, hobbies). Under the principle of network homophily, we assume that the characteristics of the communities each user is affiliated with can be inferred from aggregated information about that user’s followers, namely the self-descriptions that user’s followers provide in their Twitter biographies. For each preprint, we collected information about each user that tweeted or retweeted a link to that preprint, then queried the Twitter API to collect metadata for each of these user’s followers (**Methods**). For each user, we then compiled their followers’ biographies and screen names into a single vector of words (including emoji and hashtags), which we refer to as a “follower document.” Our subsequent analyses of these follower documents used a “bag of words” approach, that is, the order in which words occurred in each document (and the specific follower they originated from) is ignored. We refer to the collection of follower documents for a given preprint as the “audience corpus.”

For each preprint, we extracted information about the underlying audience sectors—or “topics”—by applying Latent Dirichlet Allocation (LDA) to the audience corpus of associated follower documents (**Methods**). The LDA model assumes that each audience corpus consists of *K* distinct underlying audience sectors, such that each follower document can be represented probabilistically as a mixture of these *K* audience sectors, with each audience sector represented by a collection of words that frequently co-occur within the same latent topic. We expect that each of these representations conveys semantic meaning about the characteristics of a given audience sector, e.g., a topic represented by the keywords “professor,” “university,” “genetics,” “population,” and “evolution” likely reflects an audience sector of population/evolutionary geneticists.

### Inferring the diversity of academic audiences on social media

Examples of the LDA topic modeling results for four selected preprints are shown in **Fig. 1**, where each user that referenced a given preprint is displayed as a stack of bars segmented into *K* colors representing the estimated membership fraction in each audience sector, analogous to the de facto standard of visualizing inferred genetic ancestry of individuals with the popular *STRUCTURE* program, which uses the same underlying Latent Dirichlet Allocation model [38,39]. We selected these preprints because they represent a variety of scientific disciplines covered on bioRxiv (animal behavior, synthetic biology, genomics, and neuroscience) and their audience sectors exhibit a wide range of academic and non-academic communities. For each preprint, we classified each of the inferred audience sectors as “academic” or “non-academic,” flagging topics whose top keywords include a combination of words indicating an academic affiliation, such as “PhD,” “university” or “professor” as “academic audience sectors” and classifying all other topics as “non-academic audience sectors.” For the academic audience sectors, we took the additional step of attempting to map the words associated with each topic to a singular scientific discipline (**Methods**).

**Fig. 1.**
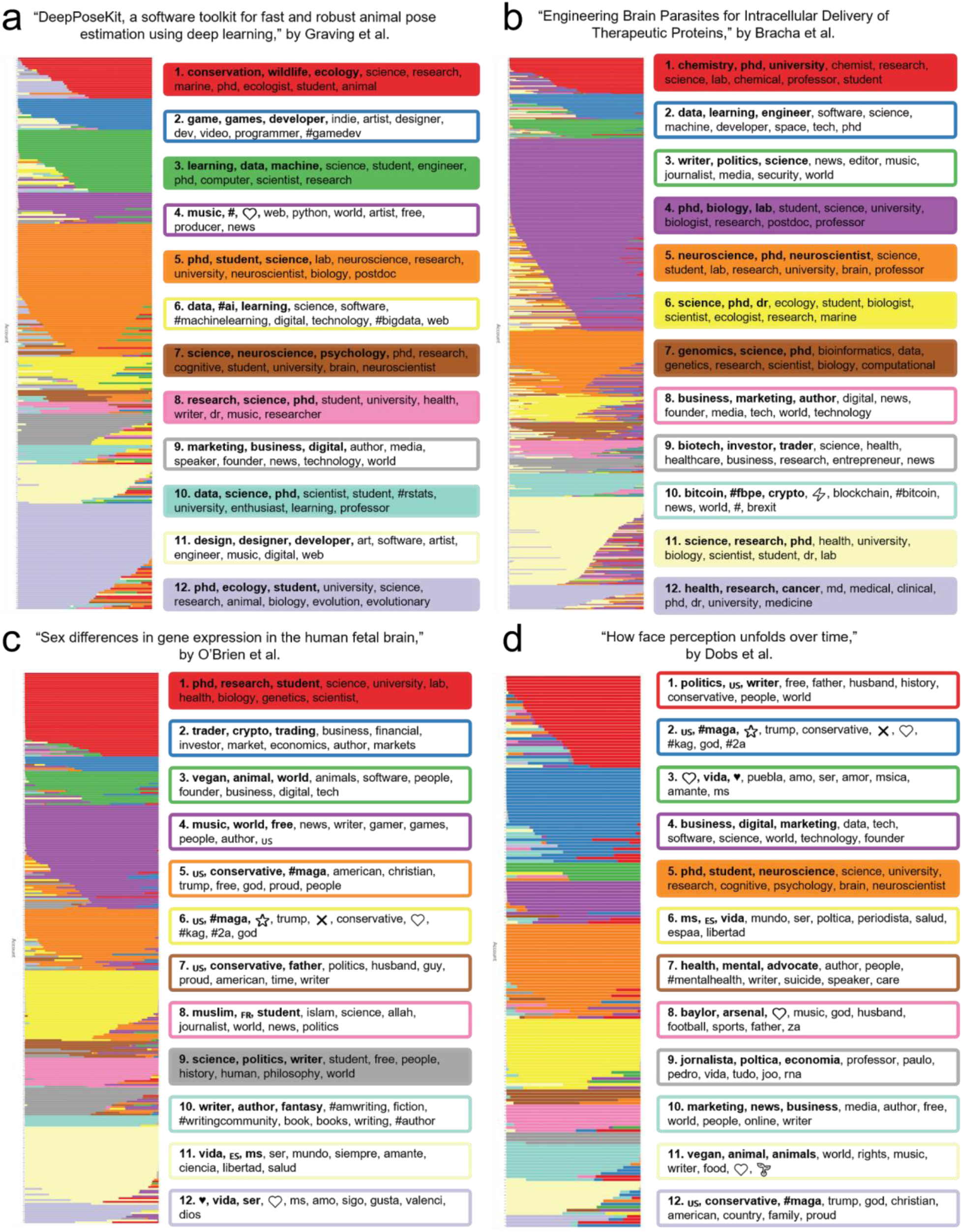
Topic modeling output for 4 selected preprints. Each panel shows the LDA topic modeling results for one of four selected preprints: **A.** “DeepPoseKit, a software toolkit for fast and robust animal pose estimation using deep learning,” by Graving et al. **B.** “Engineering Brain Parasites for Intracellular Delivery of Therapeutic Proteins,” by Bracha et al. **C.** “Sex differences in gene expression in the human fetal brain,” by O’Brien et al. **D.** “How face perception unfolds over time,” by Dobs et al. In each panel, every account that tweeted about that paper is represented by a horizontal stack of 12 colored segments that represent the account’s estimated membership fractions in each of the 12 inferred audience sectors. The top 10 keywords, hashtags, or emoji associated with each topic are shown in the corresponding colored boxes next to each panel, with the top 3 keywords for each topic highlighted in bold. Topics inferred to correspond to academic audience sectors are displayed in shaded boxes.

As shown in **Fig. 1**, the academic audience sectors frequently align with the bioRxiv category under which the preprint was classified, confirming that our method captures relevant information about the characteristics of individuals engaging with that preprint. “DeepPoseKit, a software toolkit for fast and robust animal pose estimation using deep learning,” by Graving et al., was submitted to bioRxiv in the “animal behavior and cognition” category [40]. Academic audience sectors mapped for this preprint include ecology (topic 1), artificial intelligence (topics 3 and 10), neuroscience (topic 5), psychology (topics 7 and 8) and ethology (topic 12) (**Fig. 1a**). “Engineering Brain Parasites for Intracellular Delivery of Therapeutic Proteins,” by Bracha et al., was submitted to bioRxiv in the “synthetic biology” category [41]. Academic audience sectors mapped for this preprint include molecular engineering (topic 1), bioinformatics (topic 4), neuroscience (topic 5), evolutionary sociology (topic 6), biostatistics (topic 7), public health (topic 11), and medical psychology (topic 12) (**Fig. 1b**). “Sex differences in gene expression in the human fetal brain,” by O’Brien et al., was submitted to bioRxiv in the “genomics” category [42]. Academic audience sectors mapped for this preprint include bioinformatics (topic 1) and political philosophy (topic 9) (**Fig. 1c**). “How face perception unfolds over time,” by Dobs et al., was submitted to bioRxiv in the “neuroscience” category [43]. This preprint was found to have only a single academic audience sector, psychology (topic 5) (**Fig. 1d**).

One way we can attempt to coarsely quantify the broader societal impact of a preprint using these data is to compare the relative engagement from academic versus non-academic audiences on social media [44], similar to how Altmetric provides an estimated demographic breakdown of users tweeting about a paper [21]. For each preprint, we estimated the fraction of the audience falling into any of the inferred academic audience sectors and non-academic audience sectors (**Methods**). We found that the academic audience sectors typically comprised the majority of a preprint’s audience—across the 1,800 preprints analyzed, we estimated 95.9% had a majority audience consisting of academic audience sectors (**Fig. 2**).

**Fig. 2.**
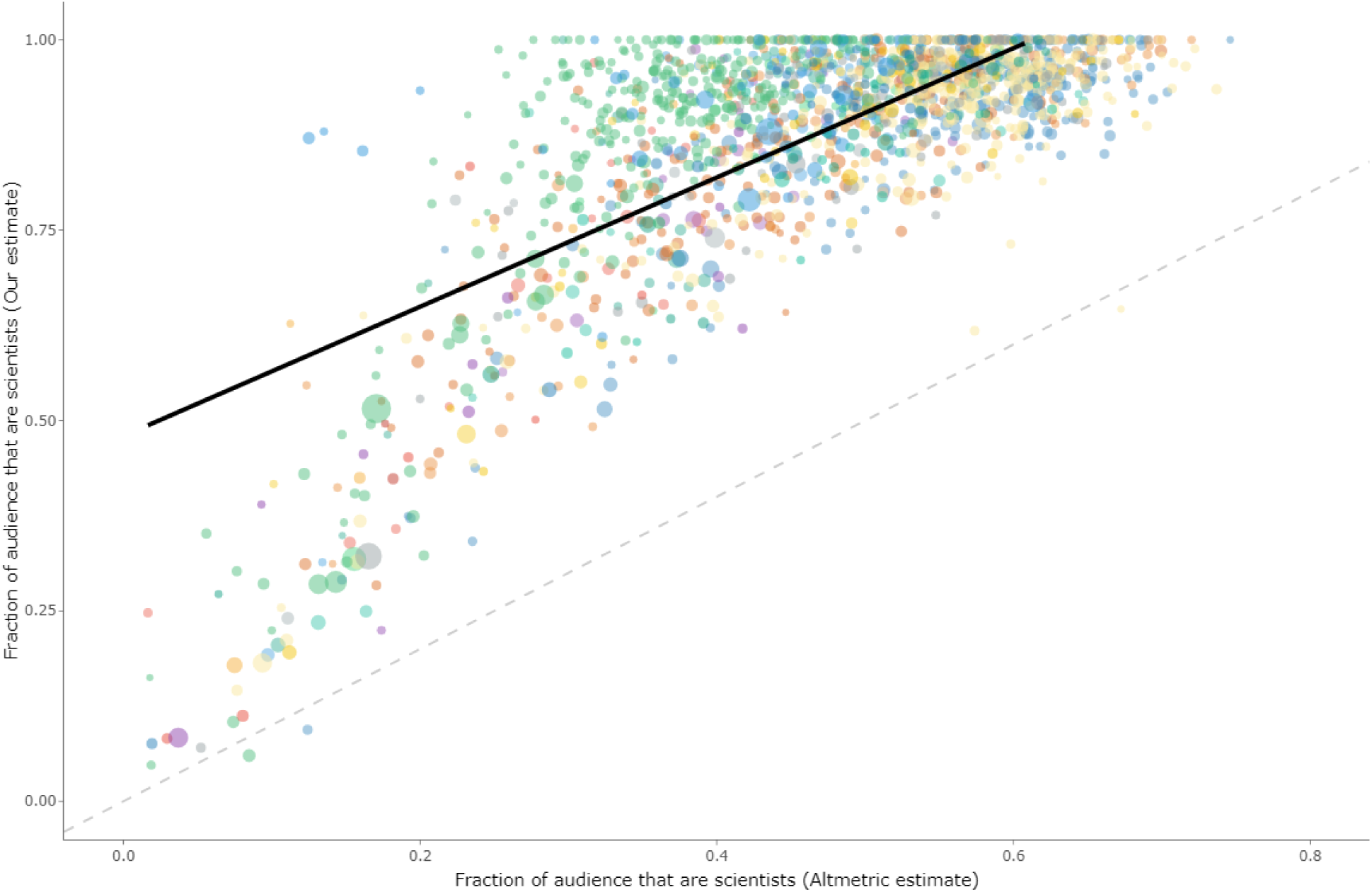
Comparison of academic audience fractions estimated by Altmetric versus our estimates. Each point represents an individual preprint, with colors corresponding to the bioRxiv category. The size of each point indicates the total number of tweets referencing that preprint. An interactive version of this figure can be accessed at https://carjed.github.io/audiences.

We next collected the demographic classifications for each preprint as estimated by Altmetric and compared these to our estimates to determine whether our estimates were consistent. The fraction of academic users for each preprint was well-correlated with the fraction of users inferred to be “scientists,” “science communicators,” or “practitioners” by Altmetric (*r* =0.72), though our method estimated a higher proportion of academic-affiliated users for all but two of the 1,800 preprints (**Fig. 2**). In many cases, a preprint’s majority audience (either academic or non-academic) as estimated by Altmetric was reversed when estimated using our method: 833 (46.3%) of the preprints analyzed had majority non-academic audiences according to Altmetric, but majority academic audiences according to our method. These conflicting estimates are difficult to reconcile, as the only way to establish the ground truth would be to manually classify each of the 331,696 users in our dataset. Though performing this manual validation is not feasible at scale, we did so for one preprint, “Shake-it-off: A simple ultrasonic cryo-EM specimen preparation device,” [45] as a proof-of-concept to demonstrate that our method is likely a more accurate classifier of academic-affiliated accounts than Altmetric’s method. Our rationale for selecting this preprint was that it received relatively few tweets that needed to be manually annotated but had high discordance between our estimated academic audience fraction (100%) and that estimated by Altmetric (39.8%). In addition, the title of this preprint contains a pithy reference to the title of the song “Shake it Off” by pop singer Taylor Swift—this was not a coincidence, as the most retweeted tweet about the preprint was posted by the first author, John Rubenstein, who described their newly-devised cryo-EM device using a parody of a line from the chorus of Swift’s “Shake if Off”:

> *Sprayer’s gonna spray spray spray…*
>
> *Shake-it-off: A simple ultrasonic cryo-EM specimen preparation device.*
>
> (https://twitter.com/RubinsteinJohn/status/112655338137443942)

This is relevant because papers with amusing, quirky or off-beat research or those that have buzzwords in the title are often presumed to attract greater attention from the public [10], so we reasoned that this preprint in particular would align with the null hypothesis that Altmetric’s estimate of a majority non-academic audience was correct for this preprint.

At the time we collected data for this preprint, Altmetric had indexed 88 unique users that had tweeted about this paper and estimated that 35 of these users were either scientists (33), practitioners (1), or science communicators (1), and the remaining 53 were members of the public. Our method only analyzed 73 of these accounts (15 were private accounts or had fewer than 5 followers and were excluded), but among these, all 73 were classified into academic audience sectors. We manually classified each of these 73 accounts as either academic or non-academic by examining self-disclosed information in each user’s bio. If a bio was empty or ambiguous, we performed a Google search for the user’s screen name and/or username to determine if they had an online presence associated with an academic institution. We determined 68 accounts belonged to individual academics (graduate students, postdocs, staff scientists, or professors), academic labs, university departments, or university-affiliated academic organizations. Further, 3 accounts (two automated bots and one human-curated account) were dedicated to posting tweets about the latest papers in the field of biophysics/microscopy and were primarily followed by academics. One other account belonged to a company that manufactures and sells microscopes and ostensibly has a customer base (and thus, Twitter following) consisting of primarily scientists. This left only a single account that we could not unambiguously classify as an academic account, though deeper examination found that many of this account’s followers work within the field of biophysics and microscopy, suggesting they also possess an affiliation with this academic network. We conclude that Altmetric misclassified over 50% of the academic (or academic-affiliated) accounts as “members of the public,” whereas our method, at worst, only misclassified a single user. This validation example demonstrates that our method of inferring audience demographics is potentially much more accurate at classifying academic-affiliated accounts than the Altmetric method.

We also note that Altmetric’s estimates of academic audiences comprised a majority for only 13 (27.1%) of the 48 preprints analyzed in the “scientific communication and education” category. Nearly all preprints we analyzed in this category revolve around topics that we interpret as being primarily relevant to academics, including several surveys of graduate students, postdoctoral fellows, and tenure-track faculty, investigations of biases in publication, peer review, and grant-funding practices, and scientometrics and altmetrics. In contrast, our method consistently estimated that academic sectors comprised a majority of the audiences for these preprints, with a median academic audience fraction of 92.6% and a minimum of 61%.

### Inferring the characteristics of non-academic audience sectors

A commonly cited advantage of altmetrics over traditional scientometrics is that altmetrics can capture information about engagement from lay audiences, which could be useful for evaluating dimensions of research productivity and scholarship that are often overlooked. For example, such information may guide researchers in describing and quantifying the broader impacts of their work in grant applications, enable new citizen science collaborations (such as fold-it, a crowdsourced computer game where players predict protein structures [46]) or crowdfunded research (such as the ∼1,000 research projects that have been successfully funded on the scientific crowdfunding platform, experiment.com), and provide concrete evidence of researchers’ individual science communication and public engagement efforts.

Overall, we observed a weak positive correlation (*r* =0.258) between the non-academic audience fraction and the total number of tweets, suggesting that preprints which gain more attention on Twitter do tend to have a broader reach with non-academic audiences. Below, we examined the non-academic audience sectors inferred by our topic modeling analysis for the four preprints used as case studies in **Fig. 1** and investigated how this information could be used in contextualized evaluation of altmetrics.

As with the inferred academic audience sectors, the characteristics of non-academic audience sectors often aligned intuitively with the topic of the preprint. “DeepPoseKit, a software toolkit for fast and robust animal pose estimation using deep learning,” by Graving et al. [40] included non-academic audience sectors associated with video game developers (topic 2), business applications of artificial intelligence (topics 6 and 9) and graphic designers (topic 11) (**Fig. 1a**), presumably because this research has implications for more realistic computational rendering of physiological properties. “Engineering Brain Parasites for Intracellular Delivery of Therapeutic Proteins,” by Bracha et al., [41]. included non-academic audience sectors primarily associated with biotechnology companies (topics 2, 8, 9, and 10), aligning with the paper’s focus on bioengineering, as well as an audience sector associated with news media (topic 3), suggesting this preprint caught the attention of science journalists (though we note that at the time of writing, Altmetric has not indexed any news media coverage which cites this preprint) (**Fig. 1b**). “Sex differences in gene expression in the human fetal brain,” by O’Brien et al. [42] had a particularly diverse array of non-academic audience sectors, including groups associated with blockchain technology and cryptocurrency (topic 2), veganism and animal rights (topic 3), video games (topic 4), right-wing politics (topics 5, 6, and 7), and science fiction & fantasy writers (topic 10). Other audience sectors captured groups of individuals that shared a common language, specifically Arabic (topic 8) and Spanish (topic 11 & 12), indicating this preprint had a culturally and geographically diverse audience (**Fig. 1c**). “How face perception unfolds over time,” by Dobs et al. [43] included non-academic audience sectors associated with right-wing politics (topics 1, 2, and 12), Spanish-speaking communities (topics 3, 6, and 9), business and marketing (topics 4 and 10), mental health professionals and advocates (topic 7), and veganism and animal rights (topic 11) (**Fig. 1d**).

### Measuring engagement from political partisans

As exemplified by our analysis of O’Brien et al. [42] and Dobs et al. [43] (**Fig. 1c-d**), many non-academic audience sectors included keywords signaling political ideology/affiliation such as “republican,” “conservative,” “democrat,” or “liberal”; hashtags such as “#MAGA” and “#KAG”, (acronyms for “make America great again” and “keep America great,” used by Donald Trump and his populist supporters throughout his campaign and presidency) and “#resist” (used primarily by democrats who critically oppose Donald Trump); and coded emoji such as 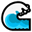 (a common symbol used by liberals to show support for a “blue wave” of Democratic candidates in the 2016 midterm elections) or **×** (used by conservatives to signal their claim that right-wing users have been unfairly subjected to censorship on social media [47]). The presence of these words, hashtags, and emoji demonstrate that many non-academic audience sectors consist of users who primarily self-identify through their political ideologies rather than professional affiliation or personal/recreational interests, and thus might be especially interested in the political implications of the research at hand.

To estimate how these audience sectors polarize on the political spectrum, we compared the fraction of users associated with topics marked with the hashtag “#resist” (an indicator of left-of-center political affiliation) to users associated with topics marked with the hashtag “#maga” (an indicator of right-of-center political affiliation) [48]. The political polarization of these audience sectors tended to vary by research area—tweets referencing preprints in ecology, immunology, scientific communication/education, systems biology, and bioinformatics came from users whose follower networks generally skewed politically left, and tweets about preprints in genetics, genomics, neuroscience, and animal behavior/cognition (and, to a lesser extent, cell biology, cancer biology, and plant biology) generally came from users whose follower networks significantly skewed to the political right (**Fig. 3**). The remaining subject areas (e.g., bioengineering, biophysics, microbiology, molecular biology) showed no significant skew in political orientation.

**Fig. 3.**
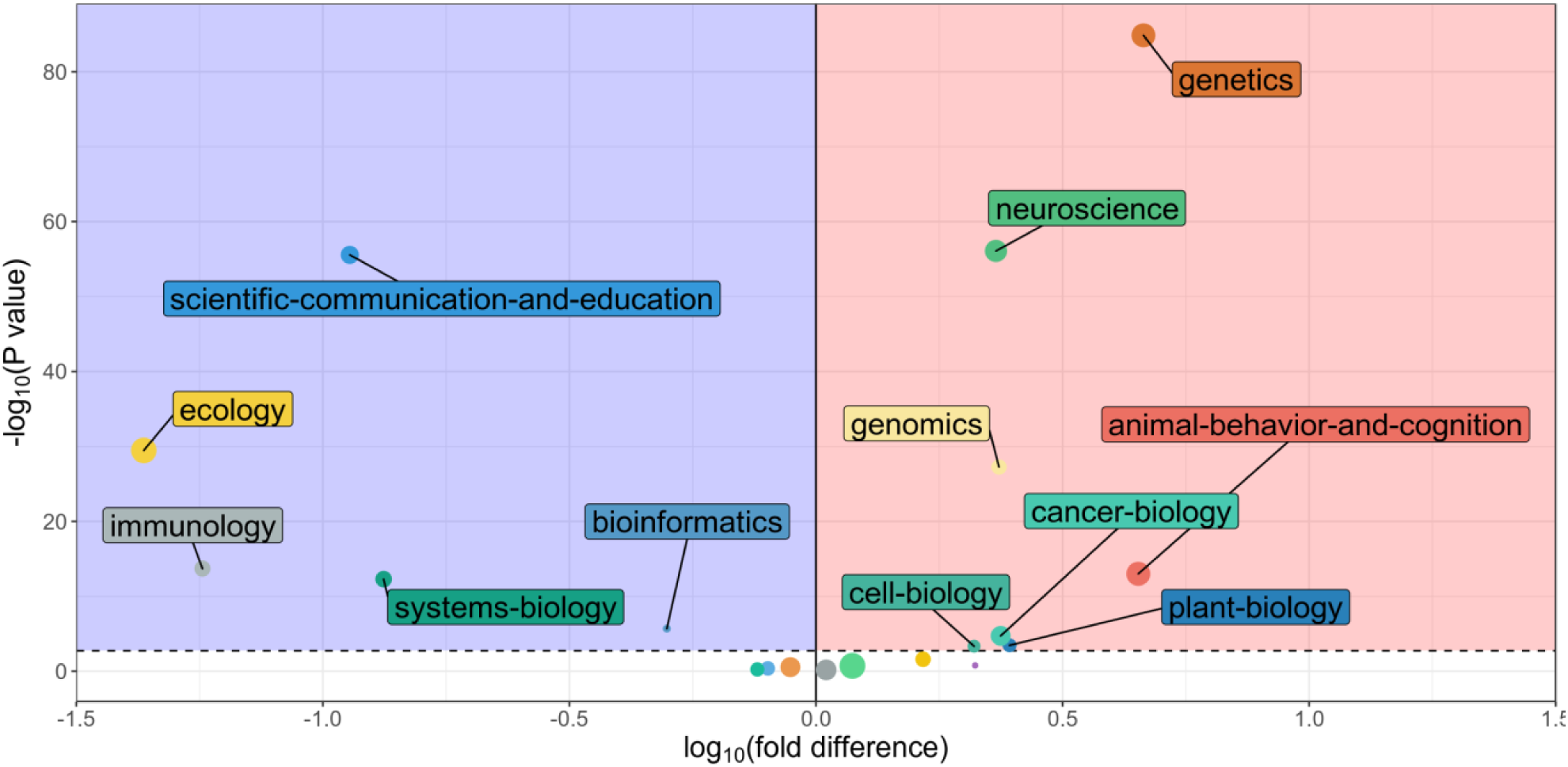
Political skew of non-academic audience sectors by bioRxiv category. The x-axis shows the log10 fold difference between the fractions of right-wing audience sectors (marked with the hashtag “#maga”) and left-wing audience sectors (marked with the hashtag “#resist”) among all tweets referencing preprints in a given bioRxiv category. The y-axis shows the -log10 p-value of an exact binomial test for whether these fractions are equal. Preprint categories with statistically significant differences (after Bonferroni multiple testing correction) are annotated above the dashed line. Preprints with audiences that skew politically left are shown in the blue shaded area, and preprints with audiences that skew politically right are shown in the red shaded area. The size of each point indicates the total number of users affiliated with political audience sectors for that category.

### Quantifying engagement with biological research by white nationalists

In addition, we found that many of the audience sectors determined to associate with right-wing political constituencies were further associated with far-right political ideologies: 1529 of the tweets analyzed were associated with audience topics that included the keyword “nationalist” in the top 50 keywords. Although users classified into these audience sectors account for only 0.24% of the 331,696 tweets analyzed, these sectors were detected in only six bioRxiv categories: animal behavior and cognition, bioinformatics, evolutionary biology, genetics, genomics, and neuroscience. This enrichment was strongest in the categories of animal behavior and cognition and genetics, where audience sectors explicitly associated with nationalism accounted for 1.53% and 1.51% of the total audiences, respectively, equivalent to a >6-fold enrichment of nationalist-associated audience sectors when compared to the overall average across all preprints analyzed.

To better quantify the extent of this engagement and confirm that this result was not an artifact of the LDA model we used, we applied an orthogonal analysis of political affiliation by examining patterns of network homophily (i.e., the number of mutual connections shared [27]) between each user that tweeted about a preprint and a curated reference panel of 20 prominent white nationalist accounts on Twitter. Across the 331,696 tweets analyzed, the median network homophily of individual users was 0.1%, with 95% of users exhibiting homophily levels of <1%. Thus, we considered any user with far-right network homophily >2% (i.e., at least a 2-fold increase in far-right network homophily compared to 95% of users in the dataset) to be affiliated with this network. We emphasize that this is a strictly quantitative affiliation, and we make no claims about the interpersonal or ideological association between an individual user and white nationalist communities on Twitter, simply that they share substantially more followers in common with particular white nationalist Twitter users than the majority of users analyzed.

We next specified varying thresholds of white nationalist network homophily (2%, 5%, 10%, and 20%) and, for each preprint, counted the number of users whose median homophily with the white nationalist reference panel exceeded each threshold (**Fig. 4; Supplementary Fig. 1**). As a point of reference, we found that 1,286 (71.4%) of the preprints analyzed had audiences where the fraction of users with >2% follower network homophily with the white nationalist reference panel was negligible (<1% of tweets referencing the preprint), indicating that engagement from white nationalist-affiliated users cannot be simply explained as a normative aspect of any research discussed on Twitter.

**Fig. 4.**
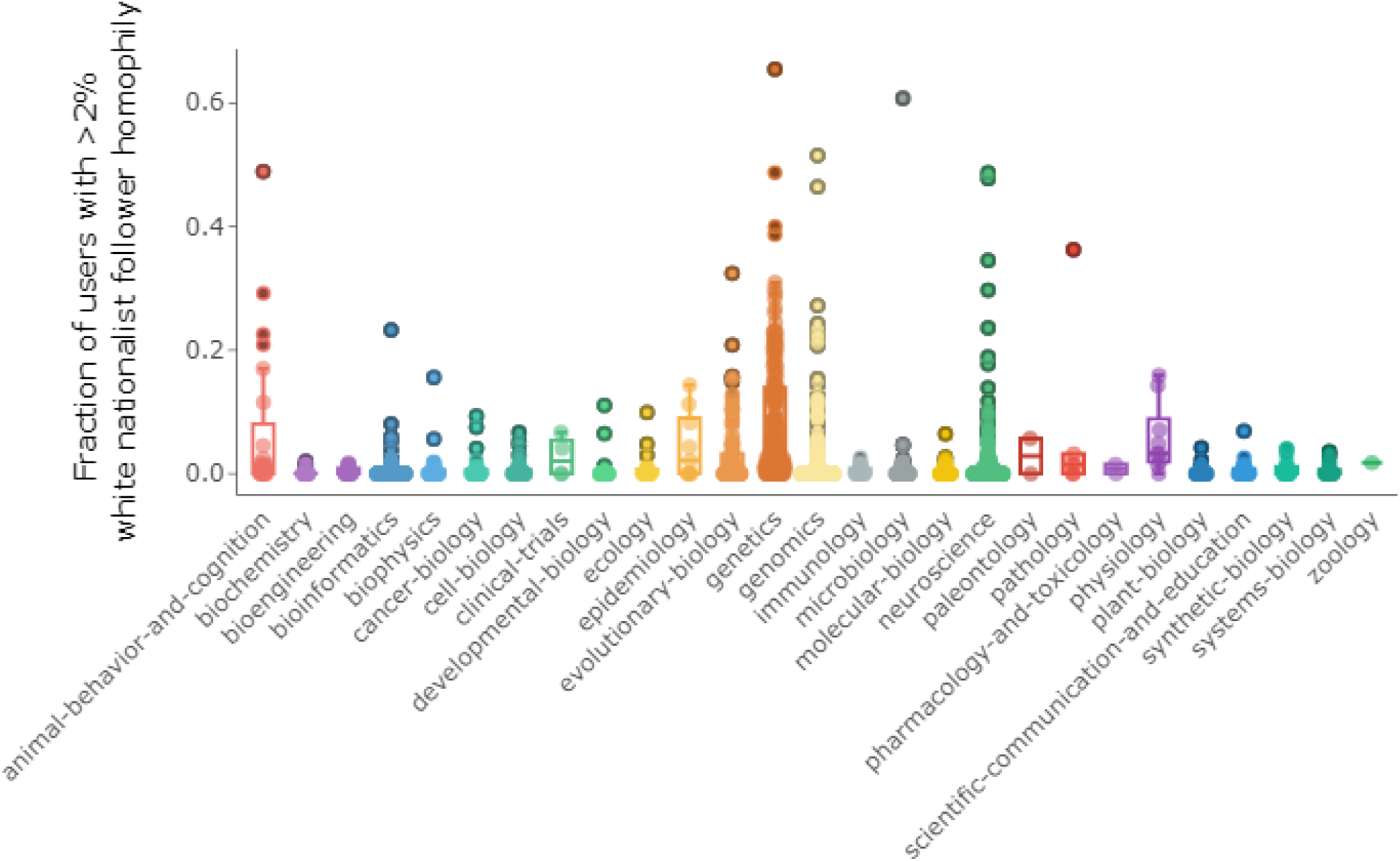
Distributions of white nationalist homophily by category. Each point represents a single preprint, and the position on the y-axis indicates the fraction of users who tweeted about that preprint whose follower network homophily with the white nationalist reference panel is greater than h=2%. Boxplots summarizing the distributions of these fractions per bioRxiv category are shown beneath each set of points.

Many of the remaining 514 preprints had an even stronger exposure to white nationalist-affiliated audience sectors than indicated by the audience topics alone: 182 of these (10.1% of all preprints analyzed) had >5% of users each with >2% white nationalist network homophily with the reference panel, and 14 of these had >30% of users with >2% white nationalist network homophily. Preprints with the strongest enrichment of such users generally occurred in bioRxiv categories where we detected explicit white nationalist associated audience sectors (animal behavior and cognition, evolutionary biology, genetics, genomics, and neuroscience), though we note that we also detected at least one preprint where >5% of tweets came from users with >2% white nationalist network homophily in 19 of the 27 categories (**Fig. 4**).

On average, the 182 preprints that showed a high level of engagement from white nationalist-affiliated accounts (>5% of users, each with >2% far-right network homophily), received 215 tweets; this is significantly more than the 1,619 preprints that did not exceed this threshold, which received 164 tweets on average (t-test, p=0.012). These 182 preprints also had significantly higher Altmetric Attention Scores (mean Attention Score of 181 compared to 112 for the remaining preprints; t-test, p=0.001), and the correlation between estimated non-academic lay audience fraction and total number of tweets was higher for these 182 preprints (r=0.38 versus r=0.22). This suggests that preprints attracting white nationalist-affiliated communities tend to receive more tweets, higher priority in Altmetric rankings, and proportionally less engagement from academic audiences.

## Discussion

Our analyses demonstrate that the audiences engaging with scientific research on Twitter can be accurately classified into granular, informative categories across both academic and non-academic communities purely by examining how each user’s followers self-identify in their Twitter biographies. This audience partitioning approach enables more accurate appraisal and contextualization of an article’s exposure to specialist and non-specialist audiences, reveals patterns of engagement from political partisans that may reflect immediate policy implications of contemporary research, and quantifies troubling trends of engagement with the scientific literature from users affiliated with far-right white nationalist networks.

It is important to note that the results for a given preprint represent a single temporal snapshot of its Twitter audience—additional users may discover and tweet about the preprint, follower networks are constantly shifting, and users may change their biographies, delete their tweets, deactivate their accounts, or be suspended at any time. The constant evolution of social media data may mean that the conclusions from our present study are only valid at a particular point in time, so tracking these trends longitudinally might be necessary to separate signals of ephemeral attention from evidence of long-term societal impacts. There are numerous other untapped features of social media data that could be useful for parsing characteristics of a preprint’s audience, such as a sentiment analysis of the tweets referencing the preprint, indexing of users who like or reply to tweets referencing the preprint, or the temporal patterns of how referencing tweets accumulate retweets.

Our study focused exclusively on bioRxiv preprints due to the accessibility of article-level metadata and breadth of research topics covered, but our audience segmentation strategy could be extended to evaluate the social media audiences of research appearing in peer-reviewed journals, too. For preprints that are eventually published in peer-reviewed journals, we could even compare the audience sectors of the manuscript in both its preprint and peer-reviewed forms to investigate how the peer review and editorial processes influence audience composition. *A priori*, we may expect the pre- and post-publication audiences to be quite similar, but many factors (press releases, journal prestige, major revisions made to the manuscript after peer review, open-access status of the paper, etc.) could alter the balance of stakeholders interested in a paper. An expansive survey of Twitter audiences for peer-reviewed publications could help journals identify strengths and growth opportunities for their editorial/publishing practices, assist authors in making informed decisions about where to submit and whether to preprint their work, and guide readers towards research that is most relevant to their interests.

Another source of data that is largely ignored in the altmetrics literature is “secondary engagement” events—tweets that do not link directly to a research article, but instead cite news articles, blog posts, and other online sources which reference or discuss the research. These secondary engagement events are not indexed by altmetric data brokers, but, given that most lay audiences learn about new scientific research through news media [8], it is likely that such posts are heavily enriched for non-academic audiences and thus more informative of potential societal impacts. For example, the paper “Loci associated with skin pigmentation identified in African populations” [49] received 374 tweets from 323 users as of December 2019, according to Altmetric. This study was covered by science journalists Ed Yong in The Atlantic [50] and Carl Zimmer in The New York Times [51] (in addition to over 40 other news outlets, according to Altmetric). Tweets linking to news articles posted by Yong, Zimmer and their respective publishers alone received over 700 retweets, far outnumbering the tweets that link directly to the research article.

### Scientists are the primary drivers of social media engagement

We estimate that academic-affiliated accounts comprise over half of the audience for over 95% of the bioRxiv preprints analyzed, suggesting that most social media discussion of preprints remains within the confines of the academic community. This stands in sharp contrast to Altmetric’s audience segmentation estimates, which imply that only about half of the preprints analyzed have majority academic-affiliated audiences on Twitter. Our results suggest that Altmetric tends to severely underestimate the engagement attributable to academic audiences. We speculate that, on average, Altmetric misclassifies up to 40% of the Twitter audience for bioRxiv preprints as members of the public, when these users should instead be classified as scientists, science communicators, or practitioners. We do not claim our method is a perfect classifier of academic audience sectors, but these discrepancies certainly highlight the need for greater transparency in the methodologies used by commercial altmetric data brokers and assurance that their altmetric indicators are situated in an informative, accurate context.

These results, coupled with the fact that scientists on Twitter are not immune to the effects of network homophily (a recent paper estimated that faculty members in ecology and evolutionary biology typically have a Twitter following comprised mostly [55%] of other scientists [14]), also leads us to conclude that most discussions of bioRxiv preprints on social media are often simply an online extension of the broader academic research ecosystem. This conclusion challenges a recent study that estimated less than 10% of tweets referencing scientific publications originated from accounts that conveyed curated, informed perspectives on the research, whereas most tweets appeared to originate from automated bots or accounts the authors characterized as duplicative, mechanical, or “devoid of original thought” [11] (though this study only considered papers in dentistry journals). To the contrary, our finding that scientists are the largest audience sector would suggest that the opposite is true, and most tweets referencing bioRxiv preprints represent curated, informed perspectives from subject matter experts. Although we cannot guarantee that every scientist on Twitter is immune from mechanical tweeting or performative attempts to build social capital (colloquially known as “clout” on Twitter) with their peers, we are optimistic that most are doing so in an honest attempt to stay abreast of the latest work in their fields and broadcast their informed opinions to their followers.

Nevertheless, the overwhelming ubiquity of academics in the audience for bioRxiv preprints reflects an obvious downside: most preprints ultimately appear to be receiving limited engagement from lay audiences. This presents an opportunity for motivated scientists to self-audit their own network homophily, build a following beyond their professional bubble, and use the platform for spreading scientific information to the broader public. If funding agencies and policymakers continue to prioritize societal impact and altmetrics as a desirable outcome of scientific research and demand evidence of impactful scholarship, it may be beneficial to explicitly incentivize such public engagement and reward researchers who develop and maintain a public-facing presence on social media.

Our analysis of Graving et al. provides a motivating example of how our social media audience segmentation approach can help recognize potential downstream societal impacts of research output and the efforts of individual researchers (**Fig. 1a**). This preprint introduced a software program called DeepPoseKit that uses deep learning to understand the dynamics of how animals sync and swarm together [40]. Of the 24% of Graving et al.’s audience that we classified as non-academic, certain sectors appeared to be associated with video game developers and graphic designers, perhaps indicating this research has immediate economic and cultural applications through the visual arts. Much of the total engagement (∼270 retweets) surrounding this preprint stemmed from a single tweet posted by the first author of the study, Jacob Graving, who provided a brief summary along with an animated GIF showing how the program tracks the movements of a swarm of locusts (https://twitter.com/jgraving/status/1122043261777076224). Neither the preprint itself nor Graving’s tweet alluded to any prospects of economic, cultural, environmental, or social applications, and both focused solely on the ethological aspects of their software (though we note that this study discloses that they received unrestricted funding from graphics card manufacturer Nvidia, which may partially explain why this preprint was of interest to video game developers). Our audience segmentation of this preprint demonstrates that researchers are fully capable of capturing the attention of lay audiences simply by maintaining a presence on Twitter and creatively communicating their work.

### Uncovering patterns of political engagement

Policy implications are frequently discussed as an important example of broader societal impacts that can result from scientific research. Twitter is widely used for political discourse, so naturally our analyses identified many audience sectors marked with overtly left-wing or right-wing political constituencies that were engaging with preprints. Intriguingly, the fields that attracted politically-oriented audience sectors did not typically receive equal bipartisan attention (as we might expect for research topics that have become battlegrounds of political disagreement, such as climate change [52], or fields that typically transcend political boundaries, such as translational biomedicine [53]). Instead, when politically oriented lay audiences were present, they tended to polarize very strongly towards one end of the political spectrum or the other.

Two categories of bioRxiv preprints stood out as attracting disproportionately left-leaning lay audiences: ecology and scientific communication and education. Many of the ecology preprints we analyzed dealt with aspects of climate change, a topic that receives far more positive attention and support from left-leaning political platforms [53]. Similarly, the scientific communication and education category includes several preprints that address issues of equity, diversity, and inclusion in academic environments, which are also a prominent feature of left-wing politics. Preprints in genetics, neuroscience, and animal behavior and cognition attracted a disproportionately stronger presence from right-leaning lay audiences. These preprints often involved research pertaining to human population history and the genetic and neurological architecture and evolution of sociobehavioral traits, suggesting such research is seen by these audience sectors as especially relevant to right-wing political ideologies. A cursory examination of tweets referencing these preprints indicates these right-wing lay audiences generally view this research through a positive lens. For example, a conservative political scientist with over 80,000 followers tweeted a reference to “Genetic Associations with Mathematics Tracking and Persistence in Secondary School” by Harden et al. [54] and interpreted the conclusions of the paper as follows:

> *Want an example of how PGS* [polygenic scores] *can inform policy issues? Voila.* [sic]
>
> (https://twitter.com/charlesmurray/status/1114536610266267649).

We strongly emphasize that the authors of this particular preprint (or any other, for that matter) do not necessarily endorse the interpretations of audiences that find it interesting and relevant, but this is a concrete example of how easily research can be appropriated in arguments for or against specific policies.

We also note that several preprints in a variety of other research areas piqued the interest of right-wing audience sectors associated with a popular conspiracy theory known as “QAnon” that originated on the 4chan and 8chan message boards [55]. Preprints that attracted these conspiracy theorist audience sectors included a study of the relationship between cell phone radiation and carcinogenesis in rats [56], evidence that the 2016 “sonic attacks” on US diplomats in Cuba may have actually been the calling song of the Indies short-tailed cricket [57], an epidemiological analysis of how pandemics spread [58], and a clinical trial exploring the effects of the ketosis diet on managing type 2 diabetes [59].

Our findings of politically oriented audience sectors (particularly in the field of genetics) may trace a broader trend of emerging genetic technologies and concepts being exploited for political advantage. In January, 2020, the administration of Republican president Donald Trump, who campaigned on a vehemently anti-immigration platform, enacted legislation that allows border security officials to “collect DNA from any person in CBP [Customs and Border Protection] custody who is subject to fingerprinting…including aliens as well as U.S. citizens and Lawful Permanent Residents [60]. In July, 2019, Israeli Prime Minister Benjamin Netanyahu cited an ancient DNA study out of context to claim that ethnic Jewish populations had settled in Palestine much earlier than the ancestors of modern-day Palestinians [61]. In 2018, Democratic U.S. Senator Elizabeth Warren publicly released the results of her genetic ancestry test in response to skepticism about her claims of having Native American ancestors [62]. Systematically studying how supporters and opponents of these politicians respond to such actions may lend further insight into how politically motivated sectors of the broader public either endorse or reject the politicization of genetics research.

### The white elephant in the room

Upon further investigation of the audience sectors we defined as “right-wing,” many such sectors were also represented by keywords indicative of extreme ideologies, such as white nationalism (in contrast, we found no indications of far-left ideologies [e.g., communism] pervading the audience sectors we defined as “left-wing”). The accounts affiliated with these audience sectors typically exhibited unusually high levels of network homophily with prominent white nationalists on Twitter, confirming that our audience segmentation was not simply the result of data artifacts in our topic model. Our findings are consistent with qualitative observations made by many journalists and scientists that members and affiliates of white nationalist movements are voracious consumers of scientific research and have attempted to reignite public interest in scientific racism to promote their ideology [35,36,63,64]. Within the last two years, multiple scientific societies have released statements denouncing the misappropriation of scientific research to justify racism and promote such ideologies, and have encouraged their members to actively participate in debunking science denialism and addressing the negative impacts of their research [34,65,66].

Although these news articles and society position statements shed light on the emerging trend of white nationalist misappropriation of science, they do not offer much sense of the quantitative scope of the issue. Our study provides conclusive quantitative evidence that white nationalists and adjacent communities are engaging with the scientific literature on Twitter. Not only are these communities a ubiquitous presence in the social media audience for certain research topics, but they can dominate the discourse around a particular preprint and inflate altmetric indicators. We must strongly emphasize that we do not claim any particular user found to be associated with audience topics pertaining to white nationalism and/or exhibiting higher than usual levels of network homophily with white nationalists is ideologically associated with such movements, merely that a nontrivial fraction of their Twitter followers likely are.

Naturally, these results elicit questions about how scientists should respond. According to a recent news report, many scientists who study socially and politically sensitive topics have expressed that they are reluctant to confront politicized misappropriation/misinterpretation of their work or avoid doing so because they feel incapable of successfully communicating the complexities of their research to non-expert audiences [35]. However, recent research has demonstrated that the reach of science denialism is actually amplified when subject matter experts and advocates do not intervene, but the spread of science denialism was significantly attenuated when experts and advocates responded with factual information or addressed the faulty rhetorical techniques of denialists [67].

Even so, the tactics proven to be effective at stemming the spread of science *denialism* may not translate well to the task of stopping the spread of science *misappropriation*. As noted by Panofsky and Donovan [64], white nationalists—unlike right-wing deniers of the reality of climate change and vaccine efficacy [52,68,69]—are not necessarily filtering scientific information through a denialist mindset or extreme misinterpretations, but rather by processing through racist cognition. Panofsky and Donovan go on to conclude that “challenging racists’ public understanding of science is not simply a matter of more education or nuance, but may require scientists to rethink their research paradigms and reflexively interrogate their own knowledge production” [64]. We anticipate the results of our study will motivate researchers to engage with these uncomfortable yet unavoidable challenges of scientific inquiry and communication.

## Materials and Methods

### Data collection

Using the Rxivist API [37], we collected metadata for 1,800 bioRxiv preprints, considering any preprints ranked within the top 1,000 by total number of downloads or total number of tweets that received 50 or more tweets from unique Twitter users. Note that this is not an exhaustive collection of the most highly-tweeted preprints, as preprints posted prior to February, 2017 were excluded from the rankings by tweet count (we expect that many of these older, highly-tweeted preprints were captured in the ranking of the top 1,000 preprints by download count, but there are likely some older preprints that received >50 tweets but few downloads).

For each preprint, we queried the Crossref Event Data API for all documented tweets and retweets that referenced that paper/preprint. In instances where tweets referencing a paper were not indexed in the Crossref event database, we used the *rvest* R package to scrape equivalent information from the preprint’s Altmetric page. Specifically, we collected 1) the handle of the user, 2) the timestamp at which they (re)tweeted the article, 3) the text of the (re)tweet, and 4) if the event was a retweet, the user who originally posted the tweet. Tweets from private users or users with fewer than 5 followers were excluded. We used the *tweetscores* R package [27] to query the Twitter API for the follower metadata (account handles and biographies of up to 10,000 followers per user) of each of the *N* unique users that (re)tweeted a given preprint. Due to the Twitter developer agreement, we are unable to provide the raw data used in these analyses; the R code used to scrape the data is provided at http://github.com/carjed/audiences.

### Topic modeling

For each of the *N* users that (re)tweeted a reference to a given preprint, we concatenated the biographies and screen names of their followers into a single “document,” representing a list of all the words contained in the Twitter biographies/screen names of that user’s followers. We cleaned each document to remove punctuation and common stopwords (e.g., “a”, “the”, “is”) from 13 languages using the *tm* R package [70]. We translated emoji that occurred in the follower documents into a unique single word string according to the official emoji shortcode, taken from https://emojipedia.org, and prepended with the string “emoji” to create an alphanumeric word, e.g., the microscope emoji 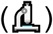 was translated to “emojimicroscope”. Similarly, we translated hashtags to replace the “#” symbol with “hashtag”, e.g.,”#microscope” was translated to “hashtagmicroscope.” For the *W* unique words observed across all the *N* follower documents, we then generated an *NxW* document term matrix, enumerating the frequencies of each word in each follower document.

We used the *lda* R package to apply a Latent Dirichlet Allocation (LDA) model to this document term matrix, representing each of the *N* documents as a mixture of *K* discrete topics (where *K*≪*N*). For consistency, we specified K=12 for all preprints. The LDA model estimates two sets of relevant hyperparameters: *θ*_*i=1,…,N,k=1,…,K*,_ indicating the probability of topic *k* occurring in the follower document for user *i*, and *ϕ*_*w=1,…,W,k=1,…K*_, indicating the probability of word *w* occurring in topic *k*. Each topic is thus characterized by words that frequently co-occur, and each follower document is summarized as a set of dosages corresponding to the probabilistic topic assignments.

### Estimation of academic audience fractions

For a given preprint, we estimated the fraction of the audience that were academics by first flagging topics containing keywords we determined to correspond to academic careers/environments (e.g., “university”, “phd”, “postdoc”, “professor”, “fellow”). We then summed the theta parameters of these topics to arrive at our estimate. Formally, this is stated as:

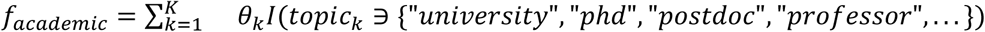

The fraction of the audience assumed to be non-academic is thus:

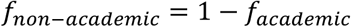

### Mapping inferred academic audience sectors to specific academic disciplines

For each audience sector inferred to correspond to an academic audience, we calculated cosine similarity scores between that audience sector’s top 50 associated keywords (*ϕ*_*w,k*_) and the 50 most frequent words found in each of the Wikipedia articles for over 1,000 academic disciplines, indexed at https://en.wikipedia.org/wiki/Outline_of_academic_disciplines. The title of the Wikipedia article found to have the highest similarity score was assigned as the best-mapping discipline of that academic audience sector.

### Network homophily analysis

We curated a reference set of 20 organizations and individuals associated with far-right white nationalist ideologies. For each of these accounts, we then scraped their list of followers. Then for each preprint analyzed, we calculated the fraction of each referencing user’s followers that were also following each of these 20 accounts, taking the median of these 20 scores as an estimate of that user’s network homophily with the reference panel. We summarized these individual-level homophily scores on a per-preprint basis by calculating the fraction of individuals with homophily score >*h*, varying *h* at four different stringency thresholds {2%, 5%, 10%, and 20%}.

## Acknowledgments

We thank Emma Beyers-Carlson, Graham Coop, Doc Edge, Jeffrey Ross-Ibarra, and Josh Schraiber for their helpful feedback on the manuscript and online resources.

