## Supplementary Information for "Quantifying and contextualizing the impact of bioRxiv preprints through automated social media audience segmentation"

### Supplementary Figures

***a.***

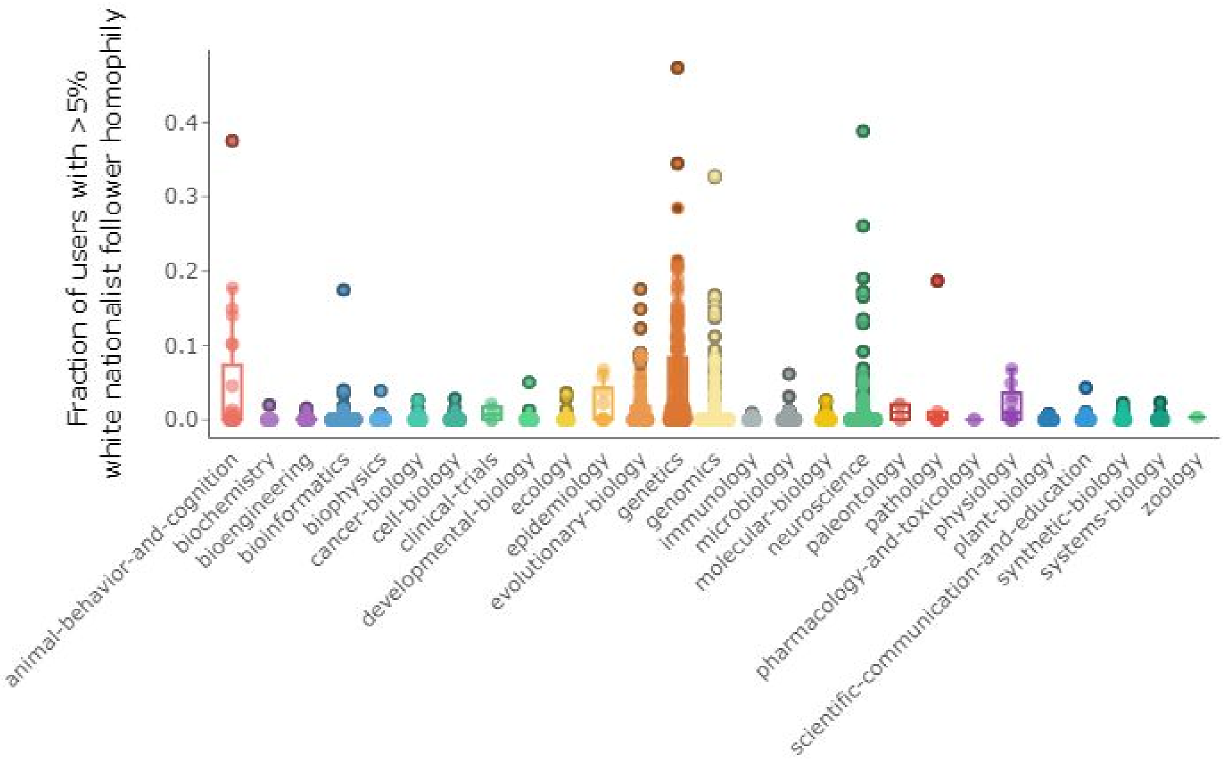


***b.***

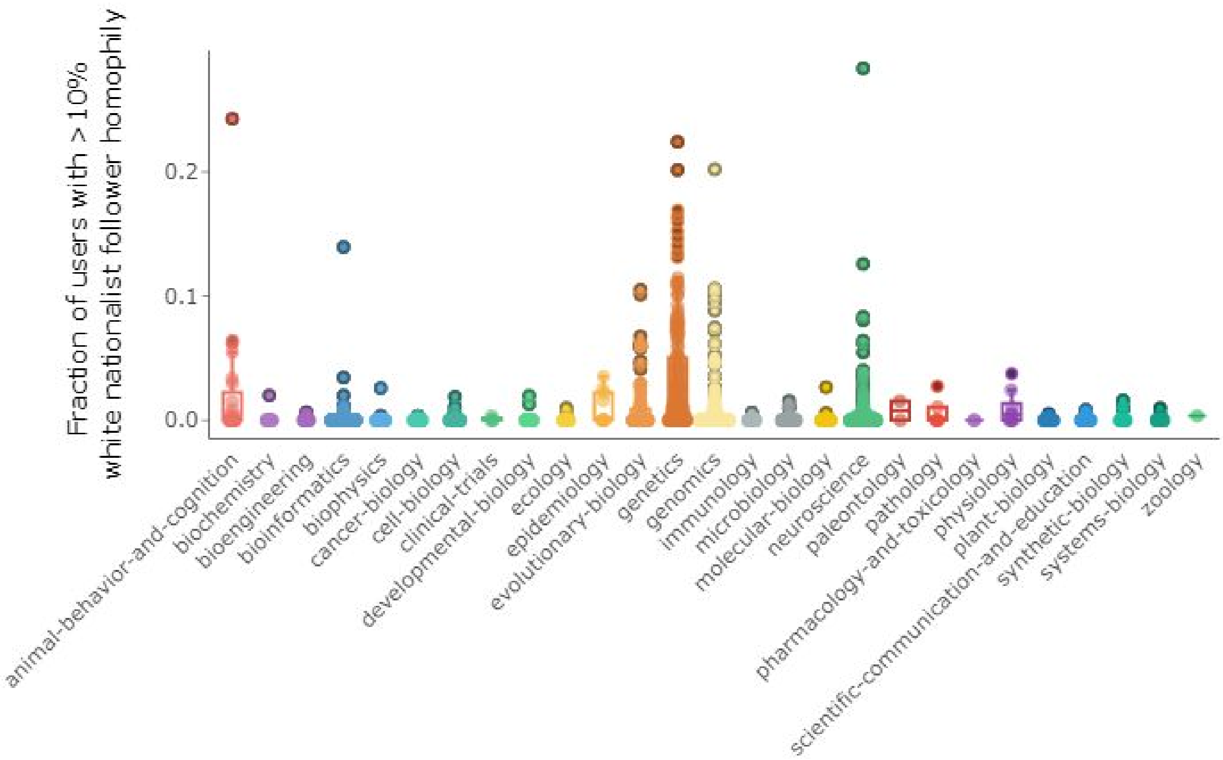
Fraction of users with >10%
white nationalist follower homophily
0.2
0.1
0.0
animal-behavior-and-cognition
biochemistry
bioengineering
bioinformatics
biophysics
cancer-biology
cell-biology
clinical-trials
developmental-biology
ecology
epidemiology
evolutionary-biology
genetics
genomics
immunology
microbiology
molecular-biology
neuroscience
paleontology
pathology
pharmacology-and-toxicology
physiology
plant-biology
scientific-communication-and-education
synthetic-biology
systems-biology
zoology

***c.***

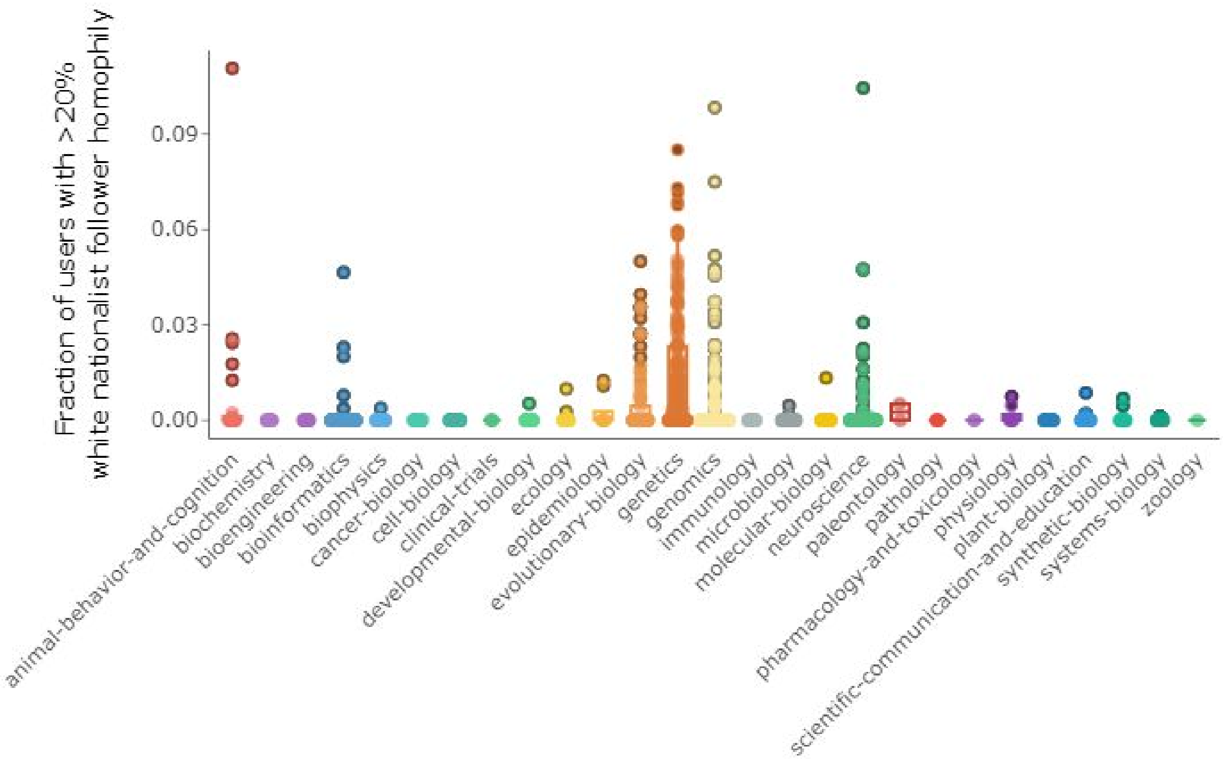


***Supplementary Fig. 1. Distributions of white nationalist homophily by category.*** *Each point represents a single preprint, and the position on the y-axis indicates the fraction of users who tweeted about that preprint whose follower network homophily with the white nationalist reference panel is greater than* ***a)*** *h=5%,* ***b)*** *h=10%, and* ***c)*** *h=20%. Boxplots summarizing the distributions of these fractions per bioRxiv category are shown beneath each set of points.*
